# Spatial patterns of biomass change across Finland in 2009–2015

**DOI:** 10.1101/2022.02.15.480479

**Authors:** Markus Haakana, Sakari Tuominen, Juha Heikkinen, Mikko Peltoniemi, Aleksi Lehtonen

## Abstract

Forest characteristics vary largely at the regional level and in smaller geographic areas in Finland. The amount of greenhouse gas emissions is related to changes in biomass and the soil type (e.g. upland soils vs. peatlands). Spatially accurate map data of forests and biomass changes could improve the ability to suggest optimal management alternatives for any patch of land, e.g. in terms of climate change mitigation. In our study, estimating and explaining spatial patterns of biomass change across Finland was the main interest. We analysed biomass changes on different soil and site types on forested land using the Finnish Multi-Source National Forest Inventory (MS-NFI) map layers of the 2009 and 2015 datasets. Silvicultural management and treatment units larger than individual pixels were created by automatic segmentation of the thematic maps. The segmented MS-NFI biomass maps of growing stocks, including above and below ground biomass resulted an average estimate of 77.7 tons ha^-1^ compared to the national forest inventory estimate of 76.5 tons ha^-1^ in 2015 for Finland. Forest soil type had a similar effect on average biomass in segmented MS-NFI and NFI data. Despite good regional and country-level results, at the segment level the biomass distributions were narrowed and averaging of biomass estimates was observed. Hence, biomass changes on segments can be considered only approximate values; also, those small differences in average biomass may accumulate when map layers from more than one time point are compared. MS-NFI classification results depend on the satellite images and field data used, causing variation in successive inventories. In addition, to avoid false biomass change observations due to the low growth rate of boreal forests, a six-year study period was set. A kappa of 0.44 was achieved for precision when comparing undisturbed and disturbed forest stands in the Global Forest Cover layer and MS-NFI segmented map, indicating the low ability of the global forest map to identify land cover changes for Finland. The segmented biomass maps provide a useful tool for forest owners to analyse carbon stock changes in their forests and how to affect the amount of carbon by forest management.

## 1 Introduction

In the EU, forest land is an important category in the LULUCF sector, absorbing nearly 9% of total emissions of other sectors in 2019 (European Environment Agency 2021). However, forest characteristics vary largely at the regional level and in smaller geographic areas. The amount of greenhouse gas emissions is related to changes in biomass and for example to the soil type (e.g. mineral soils vs. peatlands). Peatlands have huge potential in climate change mitigation (Leifeld 2018). Peatland forestry is mostly concentrated in Finland, Sweden, Norway and Russia (Strack 2008, Page and Baird 2016). It is also an important land type in the United Kingdom, Ireland, Canada, the United States and Southeast Asia. Peatland forests cover nearly 8% of forest land in the EU; the proportion is highest in the boreal zone. In Finland it is as high as 27% of which 72% is drained. Consequently, largest emissions within forest land result from drained peatlands in Finland (Statistics Finland 2021). National forest inventory (NFI) reports drainage areas for forestry land of which 4,683,000 ha have been drained for wood production according to twelfth NFI (NFI12) data (Luonnonvarakeskus 2022). Peat decomposition and related CO_2_ emissions depend on site-type where higher emissions rates are on fertile soils (Ojanen et al. 2010). Most drained forestry lands are mesotrophic and oligotrophic peatlands in Finland. Furthermore, other forest management activities such as harvesting intensity have great impact on soil carbon balance when litter input or the water table is altered.

Forest characteristics affect the forest management practices and how vulnerable single stands are to disturbances. Biomass losses, disturbances and decomposition result in greenhouse gas emissions (Herold et al. 2019). In addition to the wood production and natural values of forests, as part of forest policy, diversified forest management actions are suggested for climate change adaptation and mitigation (Ministry of Agriculture and Forestry 2015). In Finland, the forest advisor organisation Tapio publishes the guidelines for sustainable forest management, where recent findings on climate change mitigation and adaptation are incorporated. These guidelines support forest owners in their decisions and include guidance for peatland forestry. On mineral soils, forest owners can consider several harvesting options, for example selection between clear-cut-based management and continuous tree cover. Approximately 91% of productive forest land is available for wood production in Finland, i.e., potentially under intensive forest management. Hence, spatially accurate maps are valuable for forest resource management (Herold et al. 2019), and for precision forest management where decisions are made based on accurately determining forest characteristics (Holopainen et al. 2014).

During the last two decades, several maps of biomass and forest cover have been produced at global and regional scales, in many cases at coarse resolution, e.g. Hansen et al. 2003 and Kinderman et al. 2008. At a regional level, higher resolution biomass maps are also available, but in some cases, they have limitations in the available *in situ* data or the sensitivity of satellite sensors to forest biomass (Rodríguez-Veiga et al. 2016). Global forest cover change layers have been used with biomass maps to quantify biomass loss due to harvests on forest land. However, results at the European and country level show that biomass densities vary regionally, causing decreasing correlation between harvested areas and actual biomass loss. According to Ceccherini et al. (2020), biomass loss has increased by 69% between the periods of 2011–2015 and 2016–2018. This finding has been challenged and found to be conflicting with national statistics (Picard et al. 2021).

Direct field-based NFI measurements produce the most accurate reference data (Egusa et al. 2020), which can be used for the validation of biomass maps. Several countries have NFI data available, especially in the EU. However, it is often not regularly measured or up-to-date hindering biomass map validation (Avitabile and Camia 2017). In Finland, NFI data are continuously measured at a five-year inventory cycle. Besides, the Finnish Multi-Source National Forest Inventory (MS-NFI) provides multi-temporal thematic raster layers for the whole country on large number of forest variables at a pixel resolution of 16 m, e.g. timber volume and biomass by tree species, land-use class and site type. MS-NFI data is based on satellite images (Landsat 5 TM/7 ETM+/8 OLI, Sentinel 2A-MSI, IRS P6, ALOS AVNIR-2), digital maps and NFI field data and is produced every second year (Tomppo and Halme, 2004; Tomppo et al. 2008a; Mäkisara et al. 2019). In many other countries, similar products are provided (Reese et al. 2003; Gjertsen 2007; Barrett et al. 2016; Nilsson et al. 2017; Katila and Heikkinen 2020).

High resolution and reliable estimates on tree biomass and its changes are essential to policy makers to forest owners for mitigation and adaptation actions for climate change (Herold et al. 2019). Now, global climate change mitigation initiatives have connections to the forest owner level when governments support forest management practices, that increase carbon sequestration or storage in forests. In addition, the forest industry is interested in compensating for their CO_2_ emissions, where forest management actions could provide feasible ways to increase carbon sinks. Typically, forest soil is as large a carbon sink as biomass. Soil carbon modellers benefit from accurate biomass maps that they can implement smoothly in simulations.

Stands are the basic units for forest management and forestry practices prefer relatively permanent stand boundaries. Formerly, stand boundaries were delineated manually based on the interpretation of aerial photographs. Currently, they are mainly delineated automatically based on digital remote sensing data. Stands delineated from raster format thematic map data of MS-NFI based on automatic image segmentation can be used as the units for modelling of carbon cycles in forests. Pixel-based classification often has a large variation with neighbouring pixels (Yu 2006; Kim et al. 2011). In our study, we created homogeneous units by segmentation, which could be further applied in targeted climate change mitigation studies. In Finland, it is essential to delineate peatland from mineral soils due to remarkably different forest management treatments and accessibility (Pukkala 2020), and in addition, the drainage situation of peatland soils strongly affects GHG emissions. Site fertility is related to growth rate and soil GHG exchange estimation, and it is therefore important in segmentation, as well as growing stock characteristics (Mustonen et al. 2008).

Our motivation for this study was to compute a segmented biomass map according to soil type to facilitate spatial GHG exchange estimation. Biomass densities and their changes were estimated for the segments. In this study, we utilised MS-NFI data from 2009 and 2015 to derive above and below ground biomass for the segments and investigate spatio-temporal variation of biomass. At the stand the level in soil carbon modelling, it is essential to know the amount of biomass in forest stands and which kind of forests management actions have been done, e.g. allocation of felling sites, when seeking best solutions in forest management to mitigate climate change. Reference data for overall above ground and below ground biomass estimates were derived from NFI. In our study, estimating tree biomass and explaining spatial patterns across Finland were the main interests. For this we analysed biomass changes on different soil and site types on forested land (forest land and poorly productive forest land). The specific objectives of the study were (1) to produce segmented fine resolution maps on tree biomass and its changes, (2) to evaluate accuracy and usefulness of segmented biomass maps and biomass change maps, (3) to retrieve stand-replacement harvest map based on biomass changes and evaluate its precision with corresponding forest cover change map by Hansen et al. (2013).

## 2 Materials

### 2.1 Study area

The study area covers whole Finland ranging from hemiboreal to subarctic zone (59°40’-70°05’ N; 19°05’ -31°35’ E) (Fig. 1). The main tree species are Scots pine (*Pinus sylvestris*), Norway spruce (*Picea abies*), and birch species (*Betula spp*.) as well as other deciduous species (mostly *Populus tremula* and *Alnus spp*.). The forested area covers approximately 70% of the land area and is nearly entirely in the boreal zone. Topography varies typically from sea level to 200 m a.s.l., in Southern Finland, while higher hills are located in Northern Finland, where there is also a transition from forest zone to alpine tundra on the fell tops. The treeline in the north varies between 200–450 m a.s.l. (Franke et al. 2019).

**Figure 1.**
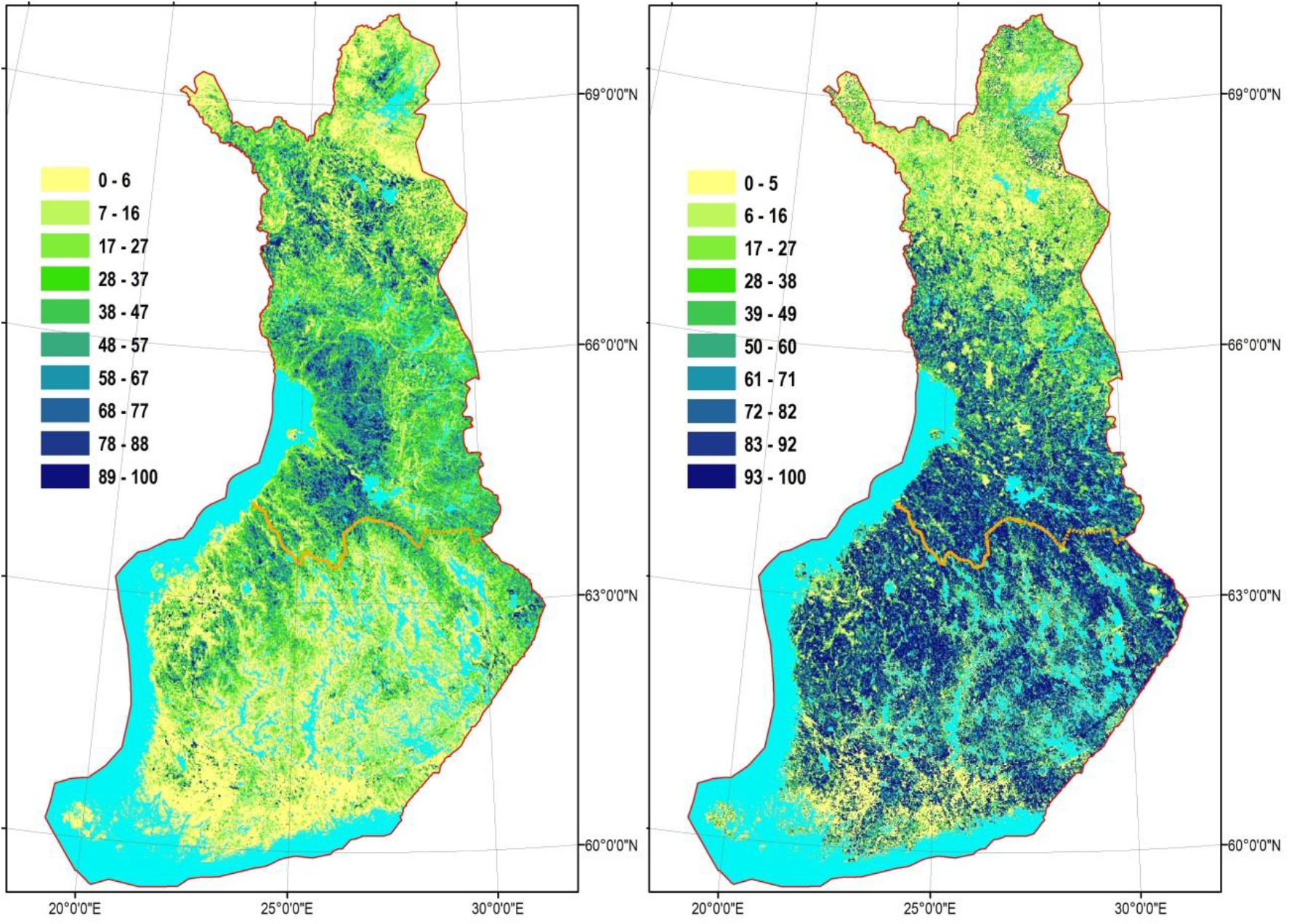
Proportion of peatlands of total forestry land (left) and proportion of drained peatlands of the total area of forestry land on peatlands (right) calculated in 1-km^2^ raster pixels. A borderline of southern and northern Finland (orange line on panels), shows the division is applied with greenhouse gas inventory data, which was also used in the Results section.

Forestry is active everywhere and on all types of forest lands where it is considered economically profitable. Exceptions are northernmost Finland and conservation areas. Forests in southern Finland consist mainly of productive land with small rocky hills or mires. Productivity decreases to the North and the proportion of peatland forests increases. Protected areas are 8.2% on forest land or 12.6% with poorly productive forest land (strictly protected forests, class 1; and protected forests on biodiversity conversation sites where cautious fellings are allowed, class 2) (Natural Resources Institute Finland 2020). Most of the protected areas are in Northern Finland.

### 2.2 Field data

National forest inventory (NFI) field data were used as a reference data in the study. NFI field plot data are based on systematic cluster sampling (Korhonen et al. 2017). We used data from the tenth NFI (NFI10 2004-2008), eleventh NFI (NFI11 2009-2012) and available data from the twelfth NFI (NFI12 2014-2017). The total number of sample plots in NFI varies between inventory cycles but is between 60 000 and 70 000 plots on land areas. NFI data are also used in the Finnish greenhouse gas inventory in biomass estimation (Statistics Finland 2021). Trees in NFI are measured whether they are sample trees or tally trees. Sample trees have larger numbers of measured parameters, and on tally trees, only basic parameters are recorded and measured, e.g. tree species and diameter at breast height (1.3 m). Biomass estimates are first predicted for sample trees by the wood density and biomass models (Repola et al. 2007, Repola 2008, 2009). Biomass estimates of tally trees are based on estimates of sample trees and the parameters of tally trees and stands (Korhonen et al. 2013 2017; 2021). In the NFI, the national definition of forest land is based on annual average growth including bark (at least 1 m^3^ ha^-1^ a^-1^); sites with growth in 0.1–1 m^3^ ha^-1^ a^-1^ are classified as poorly productive forest land. Forested land is the combination of these two classes. In this study, we also consider the FAO/FRA definition of forest land, which is applied in Finnish greenhouse gas inventory. The FAO/FRA forest land is defined as land with trees higher than five metres and a canopy cover of more than 10% or trees able to reach these thresholds *in situ* (Statistics Finland 2021). FAO/FRA forest land includes all nationally defined forest land, the most fertile part of poorly productive forest land and other areas such as forestry roads and depots.

### 2.3 Multisource national forest inventory (MS-NFI) data

We used MS-NFI raster layers covering the entirety Finland for tree biomass change estimation. The MS-NFI products are based on satellite image interpretation by non-parametric k-NN estimation method, which has been in operational use in the Finnish MS-NFI since the 1990s (Tomppo et al. 2013). The current method is described in Tomppo and Halme (2004) and Tomppo et al. (2008a, 2012). In the k-NN estimation method, NFI field plots, satellite images, digital map data and a digital elevation model are employed, and the coarse-scale variation of key forest variables are used (Tomppo et al. 2013). The resulting MS-NFI thematic maps include over 40 themes on forest parameters such as growing stock volume by tree species (m^3^ ha^-1^), site fertility class, biomass by tree species (there are four species groups) in subclasses, e.g. biomass of spruce foliage (10 kg ha^-1^). In this study, we used MS-NFI raster layers from 2009 and 2015. The MS-NFI-2009 thematic maps were based on tenth and eleventh NFI sample plot data measured in 2006–2008 and 2009–2010, respectively. The corresponding satellite images were Landsat 5 TM and for gap filling images from Landsat 7 TM, IRS P6 LISS III and ALOS AVNIR-2. The digital map data originates from the National Land Survey of Finland (NLS) and was rasterised with a 20-m pixel size in the MS-NFI (Tomppo et al. 2013). In the MS-NFI-2015, the field data originated from eleventh and twelfth NFI inventory from measurement years 2012–2013 and 2014– 2016, respectively. The NFI tree data were updated computationally by growth models to the date 31 July, 2015. The employed satellite images were Landsat 8 OLI and Sentinel-2A MSI acquired in 2015 and one scene from 2016 (Mäkisara et al. 2019). In the MS-NFI-2015 data product, satellite images and other datasets were resampled to 16 m resolution. Some areas covered by clouds were completed by the previous MS-NFI products (in MS-NFI-2015, approximately 1.2% of forestry area), in compilation sub-products from years 2000–2015 (Mäkisara et al. 2019). In MS-NFI, numerical maps of NLS were used to exclude other land classes from forestry land (e.g. arable land, roads and other built-up lands, Tomppo et al. 2013; Mäkisara et al. 2019).

## 3 Methods

### 3.1 Automatic segmentation of MS-NFI layers

Grid cell data was aggregated to larger continuous areas to resemble forest management units, which were applied in the allocation of felling sites with forest cover loss and estimation of tree biomass and its change within units. Image segmentation was used for delineating forest stands from MS-NFI thematic maps. In the delineation of homogeneous areas by their stand and soil properties, segmentation was carried out using a modified implementation of the ‘segmentation with directed trees’ algorithm (Narendra and Goldberg 1980; Pekkarinen 2002).

The segmentation method employs the local edge gradient for recognising potential segment borders. Segmentation covered forest land, poorly productive forest land and unproductive land, from which subsets of segments were selected for the study. Volumes of growing stock per main tree species (m^3^ ha^-1^) estimated in MS-NFI-2015, soil type (mineral soils, drained organic and undrained organic soils) and property borders derived from National Land Survey vector data were used as segmentation criteria. The drainage situation was computed based on NLS topographic database, where drainage was defined using a 40-m buffer around artificial ditches.

We produced segmented biomass maps and biomass change maps from MS-NFI layers as segment averages for the whole country. The class variables, like soil type and site fertility type used in the evaluation, were treated as mode variables in the segmentation. The same segment boundaries were applied for MS-NFI-2009 data

### 3.2 Validation of biomass maps

The data from NFI were used to evaluate biomass maps. We analysed biomass stocks and distribution in NFI, MS-NFI and MS-NFI segmented data. The NFI results on biomass stocks based on measurement years 2013 – 2017 were aggregated separately for southern and northern Finland from regional statistics (Luonnonvarakeskus 2019), where NFI data comprised the forested land area. Corresponding results for MS-NFI and MS-NFI-segm layers were derived from layer histograms (Table 2). Only one year NFI data, corresponding MS-NFI time points (2009 or 2015), were used for the biomass distribution analyses to avoid changes in biomass due to timber harvests or any land-use changes (Figs. 3 and 4). MS-NFI map features were extracted to the NFI plots for the analyses. Additionally, to compare the regionally summed segmented biomass stocks to the ones reported in the Finnish GHG inventory, we applied the FAO/FRA forest land definition (Fig. 5 and 6). The GHG biomass stocks were calculated separately mineral soils, undrained and drained peatland forests on FAO/FRA forest land (Statistics Finland 2021). Uncertainty associated with the NFI estimates of biomass stock was estimated as standard error due to sampling (see Tomppo et al. 2011, sec. 3.5., for details). Uncertainty due to estimation of biomass model parameters was not included, because it has no impact on validation: Both NFI and MS-NFI utilize the same models with the same errors in parameters.

Table 1 shows forest areas in NFI, GHG and MS-NFI data. NFI forested land is slightly smaller than the corresponding area in the MS-NFI-2015 layer. NFI classification is based on both land use and land cover. Therefore, in NFI, other land uses include lands with tree cover, e.g. small forest strips on built-up land. MS-NFI forest masks are land cover layers and are based on NLS digital maps. Satellite image mosaics of MS-NFI 2009 and MS-NFI 2015 are presented by Tomppo et al. (2013) and Mäkisara et al. (2019). Biomass changes were calculated for areas, where both MS-NFI 2009 and 2015 had cloud-free cover. A large part of northernmost Finland was covered by clouds; however, due to climate and altitude, only minor part of that area fulfils forest land definition. Otherwise, the remaining cloudy areas were mostly in western and northern Finland as shown in Figure 2. Average change in biomass comparisons were derived directly from cloud-free images. The change layer between MS-NFI-2009-seg and MS-NFI-2015-seg includes cloud-free areas.

**Table 1.** Forest-related areas in different datasets and total area of Finland including inland

| Datasets and different area variables | Total, 1000 ha |
| --- | --- |
| MS-NFI 2015, Total area | 33,830 |
| MS-NFI-2015, Forested land | 23,494 |
| NFI, Forested land | 22,813 |
| GHG-2009, Forest land | 21,943 |
| GHG-2015, Forest land | 21,882 |
| MS-NFI-seg change, Forested land (cloud free) | 21,371 |

**Figure 2.**
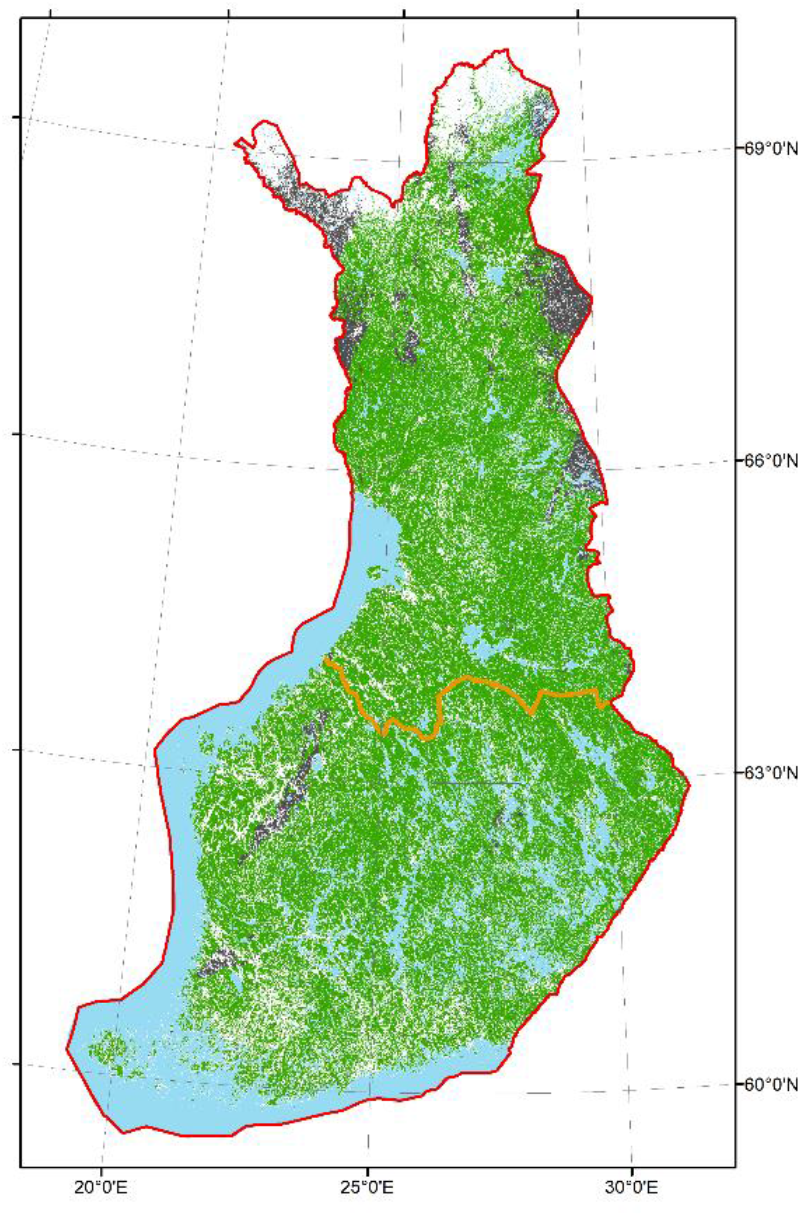
FAO Forest land (green), other lands (white), waters (blue). Clouded areas on forest land, i.e. no estimate: grey.

To evaluate the effect of biomass model errors on results, we compared also basal areas estimates between NFI, MS-NFI and MS-NFI-seg data. These results are provided in the Supplementary material.

### 3.3 Forest cover change in time-series

We selected the length of study period so that areas with biomass gains due to growth could potentially be separated from those of biomass losses due to harvests and other disturbances. The mean biomass gross increment on forest land were 3.4 t ha^-1^ a^-1^ or 20.0 t ha^-1^ within the whole study period (Statistics Finland 2021). Ideally, losses are detected in each stand when stands are harvested or affected by disturbances. Detecting biomass gains, on the other hand, may require aggregation of several stands, in time-series analyses significant trends have been found with a relatively large unit size, 1200 × 1200 m^2^ (Katila et al. 2020). In NFI11, mean volumes on mature stands on forest land were 251 m^3^ ha^-1^ and 134 m^3^ ha^-1^ in southern and northern Finland, respectively. It has been reported, for example, that MS-NFI stem volume predictions are saturated for stands over 200 m^3^ ha^-1^ (Vastaranta et al. 2014). Therefore, false losses in tree biomass derived from MS-NFI products may occur for stands with large biomass or volume.

To detect areas with stand-replacement harvests or disturbances, a threshold of 60% of mean biomass loss was used for segments.

We compared our forest loss estimates to those estimated in the Global Forest Change (GFC) dataset (Hansen et al. 2013) to evaluate the precision of that product and the conclusion given by Ceccherini et al. (2020). Global Forest Change data shows deforestation in cases where stand height has decreased below the threshold for forest land, i.e. 5 m of height. We downloaded the 2018 forest cover losses layer and applied yearly estimates from years between 2009 and 2015, version Hansen_GFC-2018-v1.6_lossyear_40N_080W.tif. Hansen’s forest cover loss map was resampled on the same grid as the segmented MS-NFI data at 16-m resolution using a nearest-neighbour method.

## 4 Results

### 4.1 Biomass estimates and distributions

Segmented and raster totals and means of biomass were slightly higher than those that were calculated from NFI plots (Table 2).

**Table 2.**
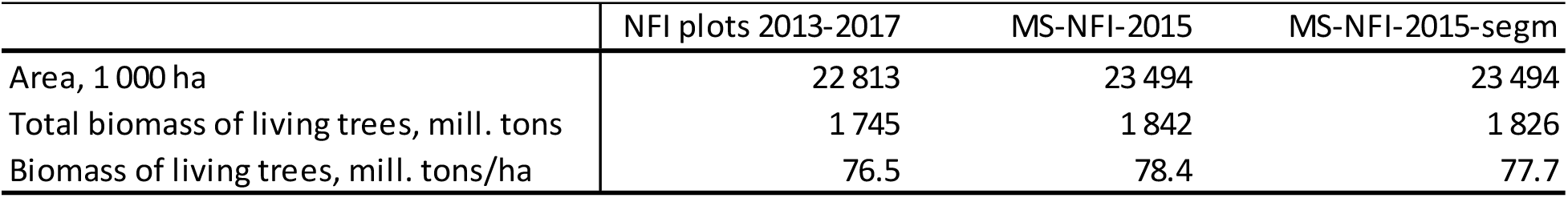
Biomass estimates for NFI plots, MS-NFI and segmented MS-NFI

|  | NFI plots 2013-2017 | MS-NFI-2015 | MS-NFI-2015-segm |
| --- | --- | --- | --- |
| Area, 1 000 ha | 22 813 | 23 494 | 23 494 |
| Total biomass of living trees, mill. tons | 1 745 | 1 842 | 1 826 |
| Biomass of living trees, mill. tons/ha | 76.5 | 78.4 | 77.7 |

NFI field data shows the greatest variation with a larger number of plots on data extremes on both inventory rounds (Fig. 3). However, MS-NFI-2009 data had very similar frequency and biomass values to NFI11. MS-NFI-2015 preserved also largely the distribution pattern as in NFI12 except for the lower end. Segmentation showed higher average biomass values at the lower end of the frequency distribution and on the other hand, the highest values were lower than in NFI. The density plot on 2009 and 2015 data shows that NFI has a relatively large number of plots with very low biomass, and plots with biomass values of more than 200 tons per hectare. Distributions MS-NFI and MS-NFI-seg data show that the largest and lowest values are averaged in the estimation process, which is the expected result (Fig. 3). The proportion of plots with over 200 tons per hectare biomasses were 4.5% and 0.2% in NFI11 and MS-NFI-2009-seg data, respectively. The corresponding proportions in NFI12 and MS-NFI-2015-seg data were 6.1% and 1.0%, respectively.

**Figure 3.**
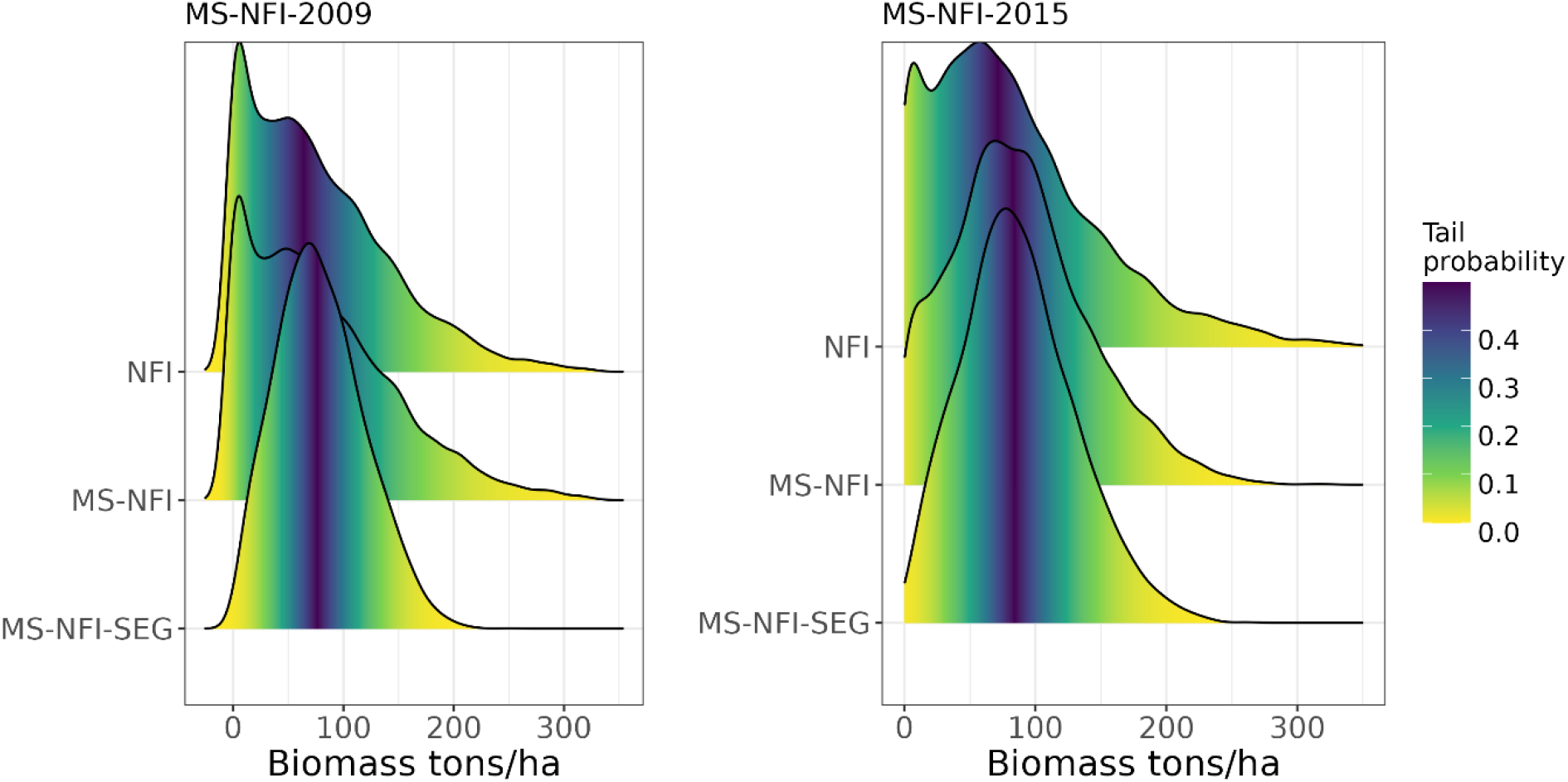
Density estimates of biomass for NFI, MS-NFI and MS-NFI-seg on forested land in 2009 (n=7590 and 2015 (n=6853) data.

Correlation diagrams of MS-NFI-2015 and MS-NFI-2015-seg data showed similar patterns of biomass stocks with NFI plot estimates (Fig. 4). In both cases correlation coefficients were of similar magnitude, but the slope of the fitted linear function was decreased in segmentation.

**Figure 4.**
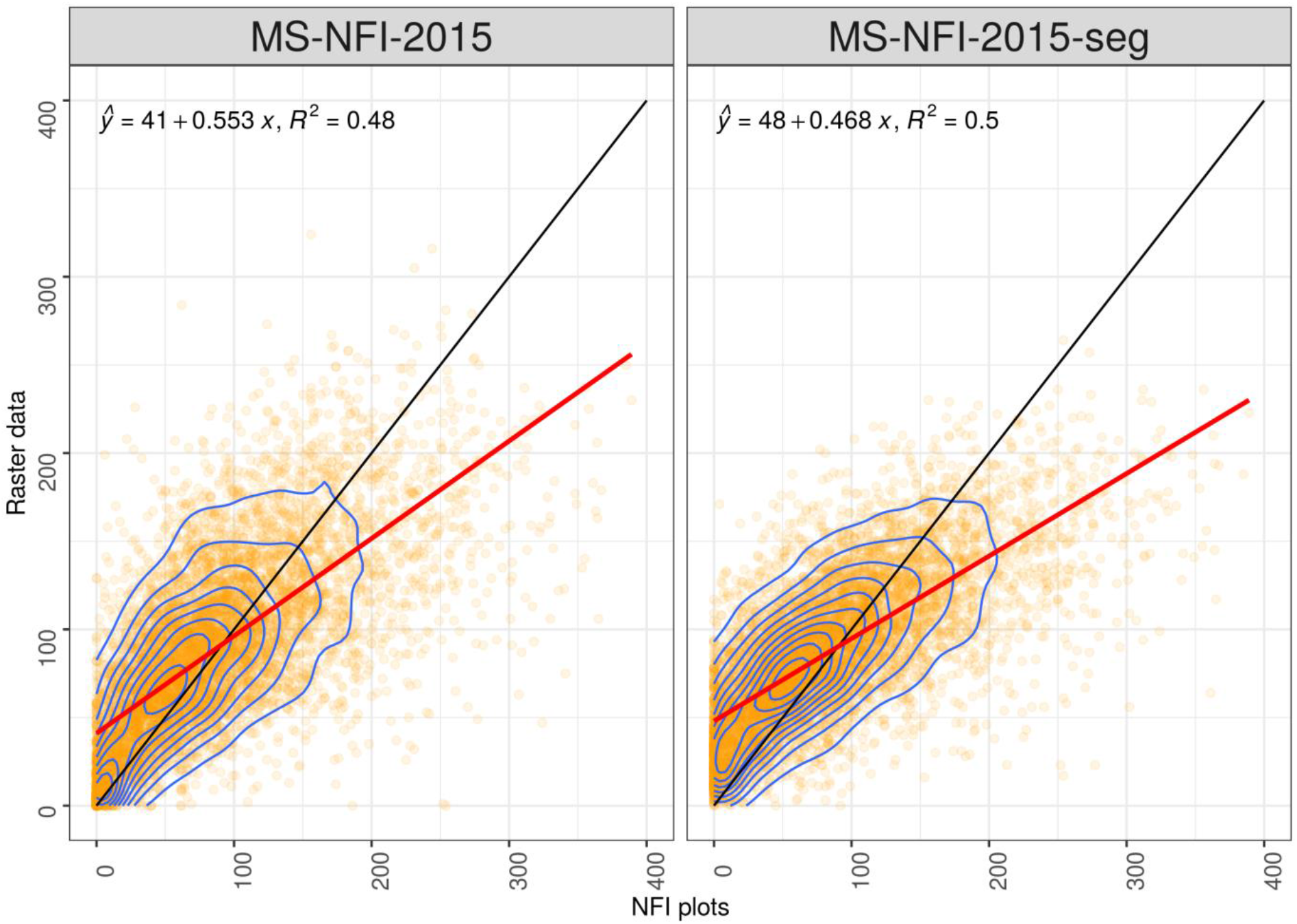
Correlation of biomass stocks (tons/ha) between NFI and pixel-level MS-NFI (left) and segmented MS-NFI (right) in 2015 data.

### 4.2 Biomass changes

Both NFI and segmented MS-NFI data showed an increase in living biomass between 2009 and 2015 (Fig 5). In 2009, NFI resulted in slightly higher biomass values than MS-NFI-seg but the average biomass values between the two datasets were still close to each other. At the same time, segmented MS-NFI showed a larger increase in biomass than NFI, i.e. 7.1 tons ha^- 1^ (+9.8%) and 4.6 tons ha^-1^ (+6.1%), respectively.

**Figure 5.**
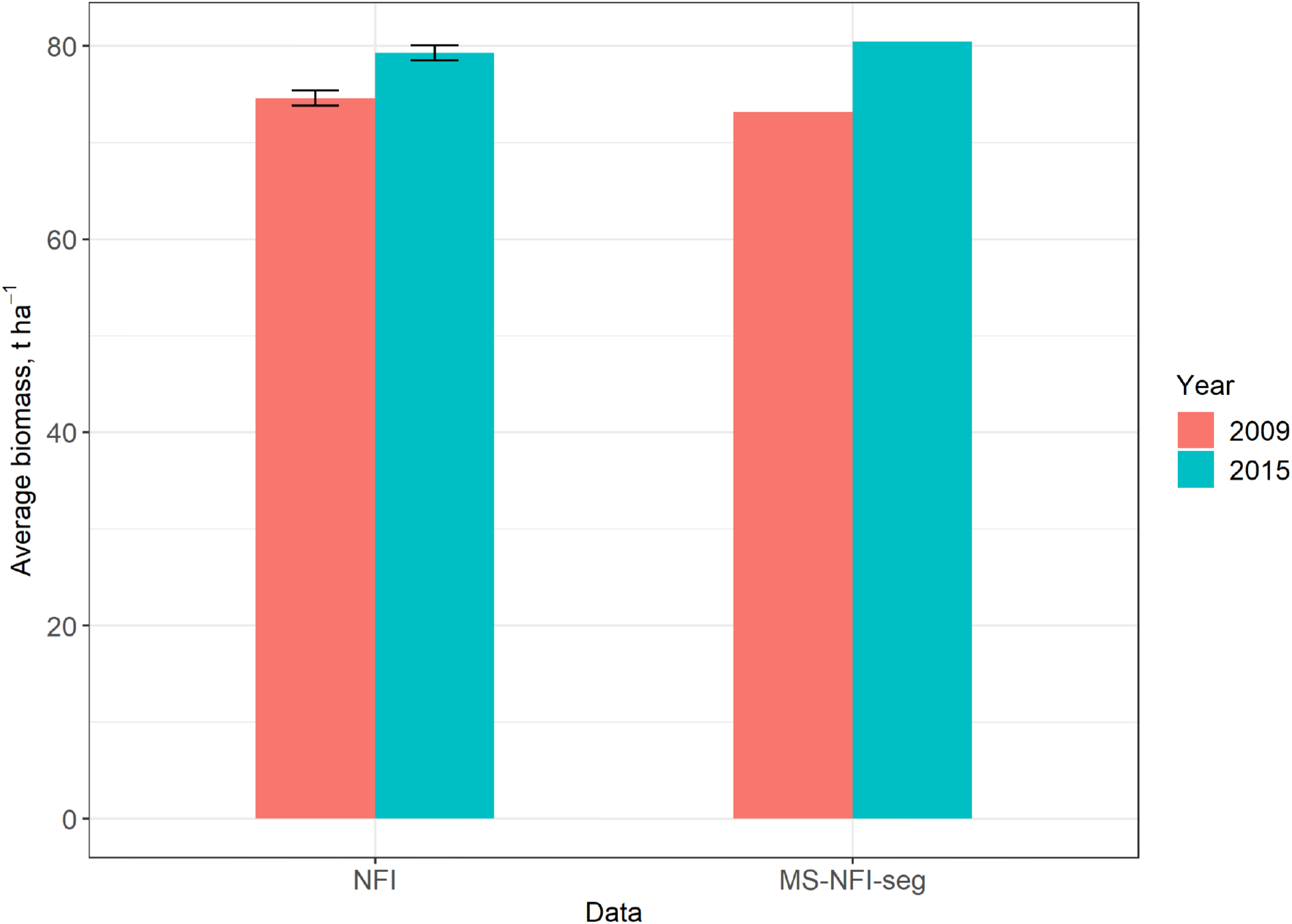
Changes in living biomass between 2009 and 2015 based on NFI field plots and segmented MS-NFI data, error bars are twice the standard error.

On mineral soils, the average biomass values were quite close to each other in NFI and MS-NFI-seg data (Fig. 6). On peatland soils, the segmented data showed slightly lower biomass values. Both data resulted in biomass increase between 2009 and 2015. However, relative changes in per hectare biomass were quite different in NFI and MS-NFI-seg in most cases except on mineral soils in Northern Finland. An exceptionally large difference between datasets was detected on undrained organic soils in Southern Finland, where NFI data resulted in a remarkably high increase in biomass compared to segmented MS-NFI data.

**Figure 6.**
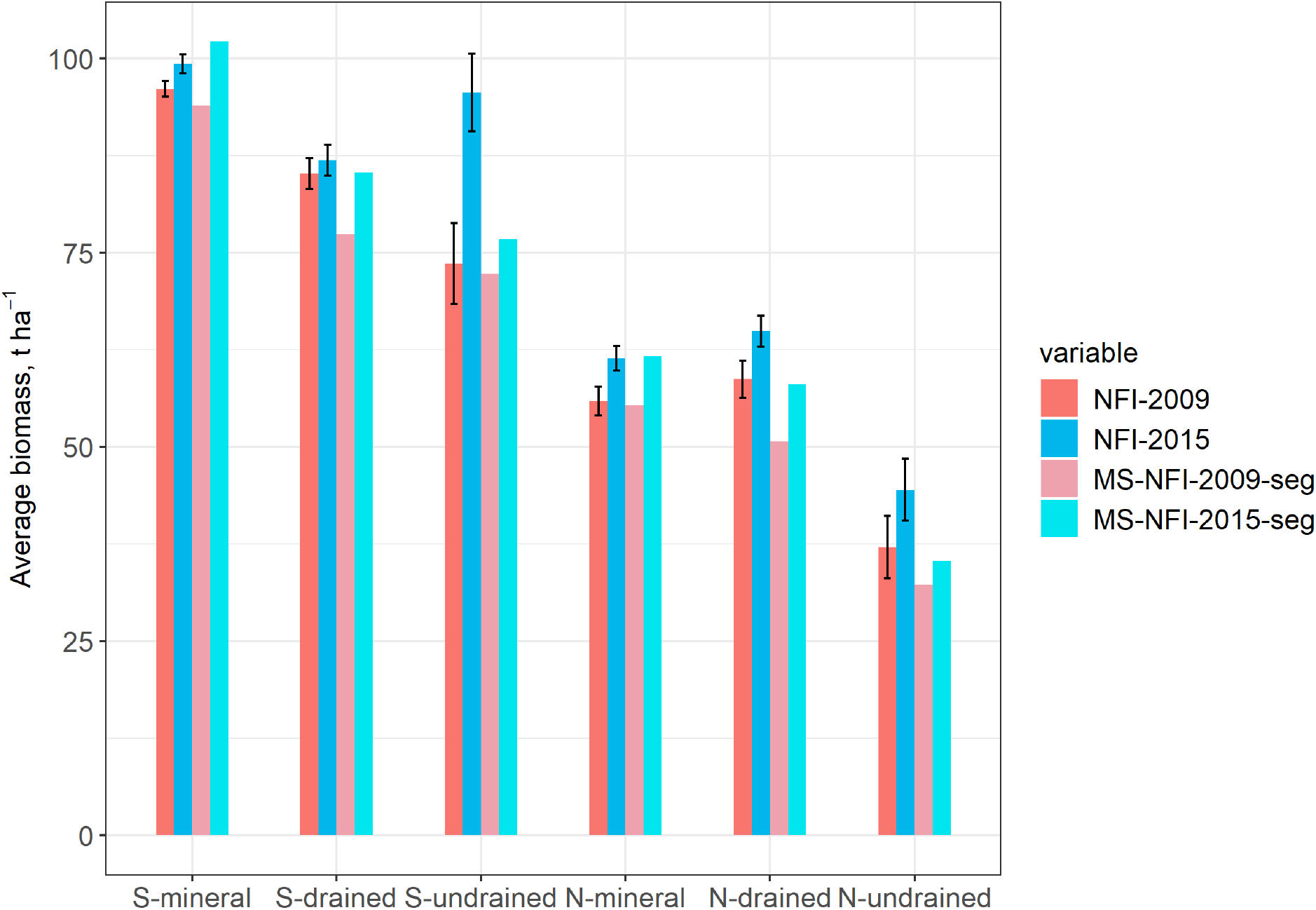
Biomass in 2009 and 2015 for different soil types in southern (S) and northern (N) Finland, error bars are twice the standard error.

### 4.3 Recognition of harvested stands

Biomass change maps showed either gains or losses for each segment and a change from forest to the non-forest stage in the case of the Global Forest Change map (Fig. 7). Forested land was covered by areas of relatively small changes in biomass, according to Luke’s forest statistics, approximately 3.9% of the forested land area was managed with stand-replacement harvests during the study period (Luonnonvarakeskus 2022). It is approximately 150,000 ha a^-1^ on average. At the segment level, the corresponding proportion was 4.9% with MS-NFI-seg data and 5.2% with the GGFC map. In the latter case, original data without segmentation resulted in a proportion of 4.2% of forest area converted to the non-forest stage. Examples in figure 7 show that for most segments, MS-NFI biomass changes were relatively small, and therefore, depending on the satellite image and field data in classification, it can be difficult to distinguish between small biomass growth and decrease. This was the case especially in mature stands with high biomass levels. However, MS-NFI-seg data includes information on biomass both before and after potential disturbance, which can be utilised for example modelling purposes together with the computed change layer. Only segments where biomass had decreased more than a threshold of 60% were interpreted as disturbed, i.e., presented with dark brown colour in figure 7.

**Figure 7.**
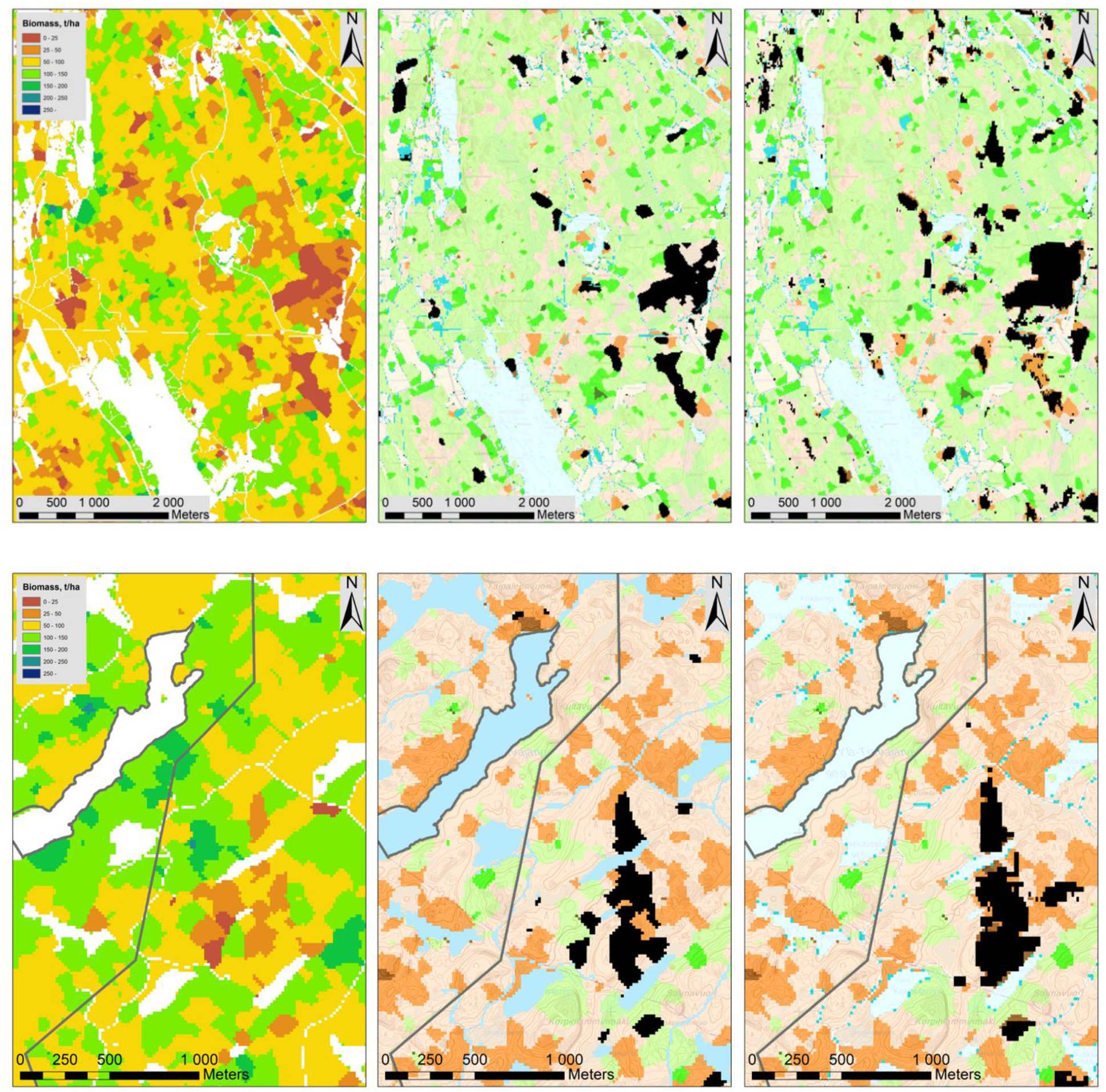
Tree biomass in MS-NFI-seg data in 2015 (left) a MS-NFI change map and deforested areas on black colour (centre) and corresponding Global Forest Change map (right) on forest cover loss between 2009–2015. Above are typical commercial forests (located in Alavus SW Finland), and below are protected and commercial forests with higher biomass (Repovesi National Park, south Finland). In the change map, the green colour indicates the amounts of gains, and the orange-brown colour shows losses. Base map is from the National Land Survey of Finland.

The confusion matrix examining the disturbed and undisturbed segments between MS-NFI-seg and the Global Forest Change map showed 95 % of correctly classified segments (Table 3). However, as the proportion of disturbed segments was very small on both data sets, the kappa value representing the success of classification was 0.44, indicating a relatively weak level of agreement between the two data sets. The producer’s accuracy of the disturbed areas was 0.40, and it was 0.98 for undisturbed areas. The user’s accuracies for the same classes were, respectively, 0.56 and 0.97.

**Table 3.**
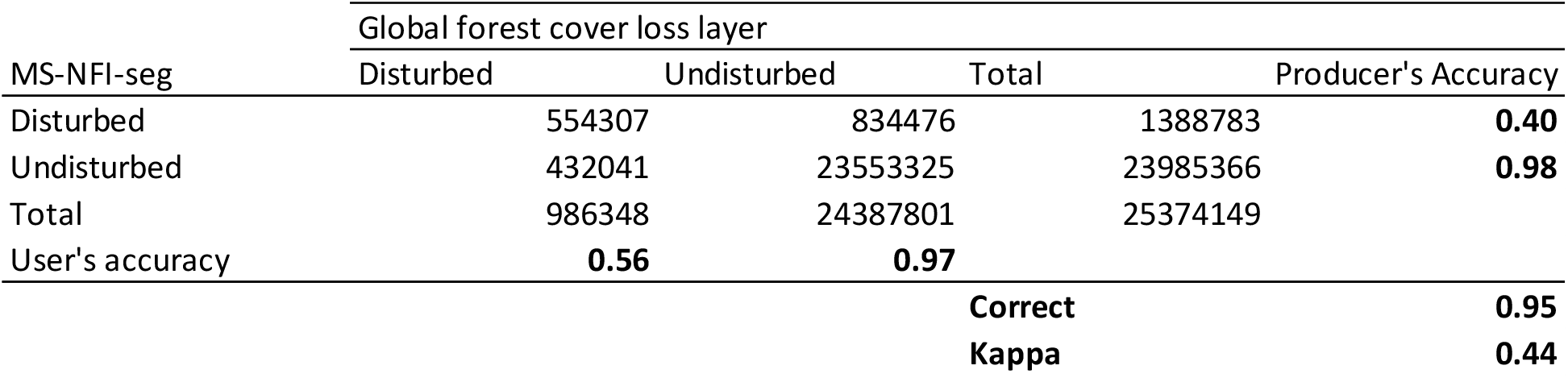
Confusion matrix of forest disturbance on segmented data set.

### 5. Discussion

The precision of biomass estimates on MS-NFI segmented data was explored in this study. This included biomass estimates at regional and whole country level, biomass change estimates and detection of harvested areas. The segmentation results were also tested with NFI field data at plot level. Especially for soil carbon modelling purposes achieving reliable estimates on segmentation is essential. Soil carbon modelling utilises information from image segments on tree biomass and their changes. Compared to pixel-level MS-NFI thematic maps, segmented data provide an effective way for modelling by speeding up the procedures. Stand properties, such as tree height, were used in delineating the segments from each other. However, also soil characteristics and property boundaries were used in running the segmentation in this study. Therefore, image segments were not homogeneous units by stand features but were affected also by management situations. In the end, stand properties did not necessarily follow segment borders, causing mixed segments.

The assessment of the MS-NFI-seg dataset on tree biomass at the whole country level showed that the estimates were very close to those derived from MS-NFI thematic maps. Compared to NFI field data MS-NFI tend to overestimate forestry land area due to it being based on digital map masks, which are land cover rather than land use. MS-NFI-2015 training data is different from NFI field data due to it being computationally updated to the target date, 31.7.2015, for which growth models were used, cuttings updated and parts of plots removed due to changes. The updating was performed separately for each processing window and calibrated with field data mean volumes (Mäkisara et al. 2019). Therefore, only minor differences between updated training data used in MS-NFI and original NFI field data were expected on biomass. In all, image segmentation did not introduce significant bias in mean and total biomass estimates compared to MS-NFI in Finland.

Segments are usually smaller than typical forest management and treatment units and reserve much of the original image variation. In MS-NFI, thematic maps are based on k-NN estimation, where the value of *k* varies from 3–5 depending on image conditions (Mäkisara et al. 2019). When a similar variation than in field data is desired, then k = 1 is appropriate, and when RMSE minimisation is desired, then a higher value of k is presumed (Franco-Lopez 2001, McRoberts et al. 2002, Katila and Tomppo 2001). K-NN-based satellite image estimates have typically high RMSEs at the individual pixel level (Tomppo et al. 2008b), which causes undesired noise in classification results when aiming at further analyses of stand variables (Hall et al. 2006). When comparing to field data, observations on data extremes tend to be underestimated when a higher number of sample records are used, which is reported in several papers, e.g. Hall et al. 2006, Tuominen et al. 2017. Even though the highest and lowest biomass values were underestimated in segmentation maps, the country-level results corresponded well with NFI. Zeng (2004) reported similar observations at regional the level. The saturation of Landsat imagery is an important factor for inaccurate aboveground biomass estimation (Zhao et al. 2016). In this study, saturation was observed on protected areas and forested land available for wood production. Protected areas included areas with relatively high biomass, even though they are commonly established in areas with less than average productivity (Häkkilä et al. 2017). Approximately 12% of forested land is protected in Finland and according to the NFI mature forests cover 12% of the total forest land area. Classification results on forested land at the mature stage are predominantly affected by pixel value saturation. However, it was noted that segmented biomass estimates are sufficiently accurate for soil carbon modelling purposes, e.g. in a way that segments can be ordered according to biomass and force cuttings in modelling to correspond statistics.

Forest areas managed for wood production have heterogeneous age structures between stands and biomass changes are caused by relatively high growth levels or a strong decrease in biomass due to timber harvesting. Image segmentation decreases slightly for RMSEs compared to pixel-level variation. However, due to the relatively short study period, RMSE at the segment level can still exceed biomass growth and results in an unreal decrease in biomass at the segment level. However, compared to the Global Forest Change map (Hansen et al. 2013), relative congruent areas were detected having biomass loss. Furthermore, the produced biomass maps provide estimates on the amounts of total biomass for soil model parametrisation. It would be also possible to combine the GFC map to MS-NFI-seg biomass estimates to get small area estimates on total biomass lost.

Harvest area estimates reported by Ceccherini et al. (2020, Extended Data Fig. 6) based also on GFC time-series and the reported average area for Finland were approximately 150,000 ha a^-1^ in the study period. However, as derived from Breidenbach et al. 2021, approximately only 100,000 ha a^-1^ was due to final harvests. The reported areas by Ceccherini et al. (2020) were quite stable during the study period but were increased between 2016–2018. Other studies suggest that this increase in the harvest area is merely an artifact (Breidenbach et al. 2021; Picard et al. 2021).

In recognition of disturbed areas, it would be better to focus on growing stock variables when segments are comprised. However, in terms of greenhouse gas emissions, soil type and management are key factors at the stand level to determine where potential sinks and sources are located. With this spatially explicit data, it is possible to provide input data for soil modellers to further study, on which forest management operations are appropriate for a certain type of stands to mitigate greenhouse emissions cost-effectively. This information can be applied to real forest management units, compartments, where similar information on soil and stand properties is available. The most reliable change estimates on biomass for segments were achieved on clear-cut sites. In case of smaller changes in biomass or tree volumes, time series analyses would improve the reliability of the change estimates (Katila et al. 2020).

## Acknowledgements

The authors want to thank the staff at the Natural Resources Institute Finland (Luke) for their contribution in the NFI data collection and preparation, and producing the MS-NFI thematic maps used in this study, and we also thank the greenhouse gas inventory project for their data. The study was funded by the Strategic Research Council at the Academy of Finland, Decision no. 312912 (SOMPA).

## Notes

### Competing Interest Statement

The authors have declared no competing interest.

