## Supplementary material for "Spatial patterns of biomass change across Finland in 2009–2015"

*Distribution of basal area*

The distribution of basal areas was tested with NFI sample plots measured in 2014–2016 and compared to MS-NFI-2015 basal area estimates and segmented data. Values of basal area are not dependent on models but are highly correlated with biomass and are used for biomass models through tree diameter (Repola 2009). In NFI, basal area is measured as an average within a stand, but for this study, also calculated basal area was produced from measured trees in sample plots. In the segmentation, stand-level basal areas were further averaged from MS-NFI classification results. Only field plots with a single stand and plots measured within one year from MS-NFI-2015 data were used for comparison. Basal areas were analysed on different soil and site types and by drainage situation.

*Basal area results*

On mineral soils and well-drained soils, there was a large proportion of stands with a high overall basal area up to 50 $m^2$ $ha^{-1}$ in NFI (Fig. S1). In MS-NFI, data maximum basal areas were slightly over 30 $m^2$ $ha^{-1}$, but in segmented data that was only true in some cases. It can also be noted that segmented values were concentrated more around the average of basal areas on mineral and well-drained organic soils. On undrained and poorly drained organic soils, NFI data and segmented data had quite similar patterns, where peak values and a load of distribution were at the lower end of frequency. However, NFI data had higher densities than MS-NFI data or especially segmented data at the lower end of the basal area distribution.

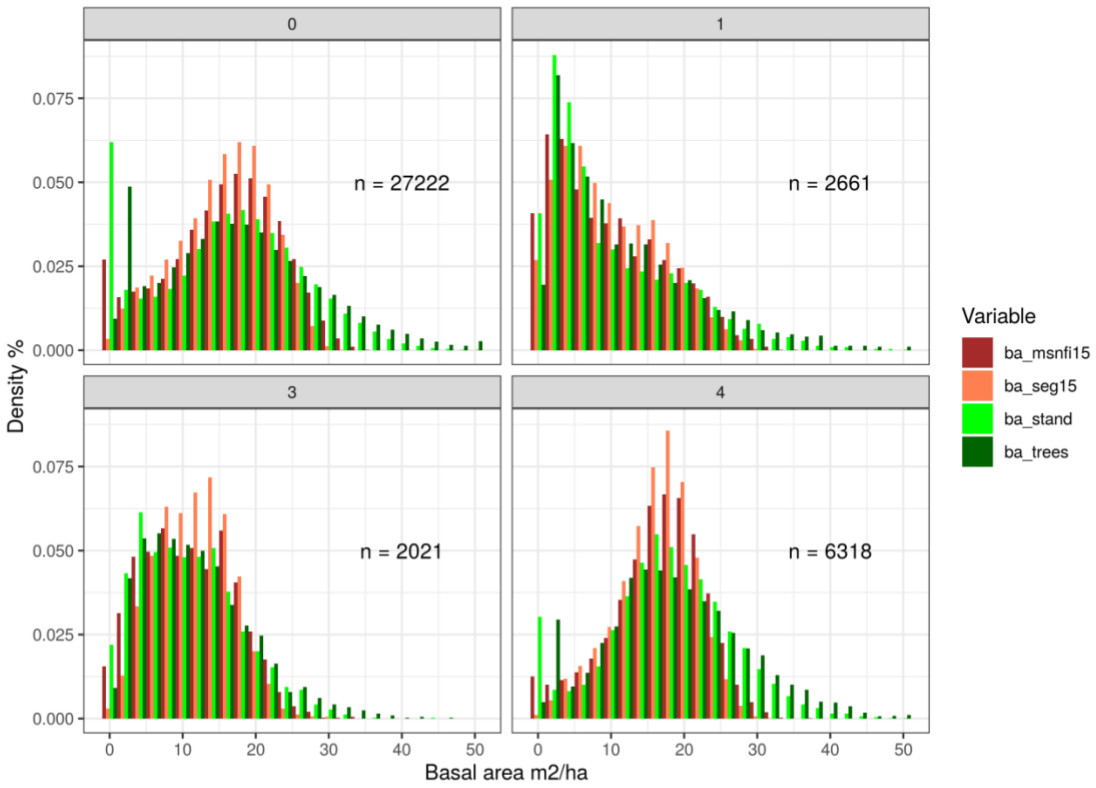


**Figure S1.** Basal area for trees on NFI plot, stand around a plot, MS-NFI and segmented data grouped by drainage situation. Mineral land (0), organic soils: undrained (1), recently or poorly drained (3), well-drained (4).

Regarding different site types, segmentation had more similar distributions with NFI on less fertile sites where maximum basal areas were relatively low (Fig. S2). On fertile sites, NFI showed higher densities at both ends of the basal area distribution. A major part of the frequency is, however, detected by both MS-NFI and segmentation except the highest basal areas.

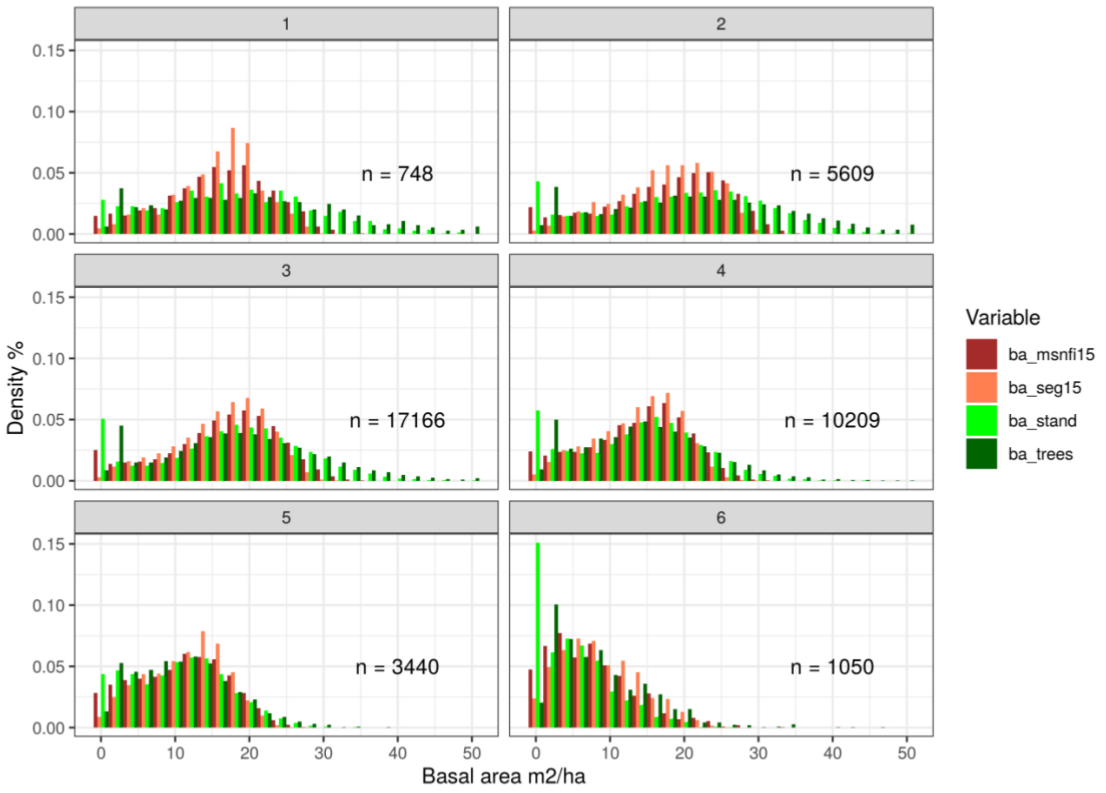


**Figure S2.** Basal area for trees on NFI plot, stand around a plot, MS-NFI and segmented data grouped by site type. Site types are at an ordinal scale from fertile (1 and 2) to poor soils (6).

Comparison according to protection status of forests showed larger mean values for commercial forests than for protected areas (Fig. S3). The largest part of the protected forests is located in northern Finland on less fertile lands where mean biomass values are lower. Smaller areas, especially in southern Finland nature reserves, also include fertile lands with higher mean biomass values. Regardless of the protection status of the forested area, segmentation produced congruent results and showed that the model behaves the same way in these management situations. The standard deviation of basal area in NFI data was much larger than in MS-NFI or MS-NFI-seg data, indicating that the smallest and highest values of basal areas were underestimated in segmented MS-NFI. On the other hand, average basal areas of MS-NFI-seg corresponded closely to NFI observations at the stand level in both protected and commercial forests. Standard deviation was slightly higher in protected areas for segmented data and NFI than in commercial forests.

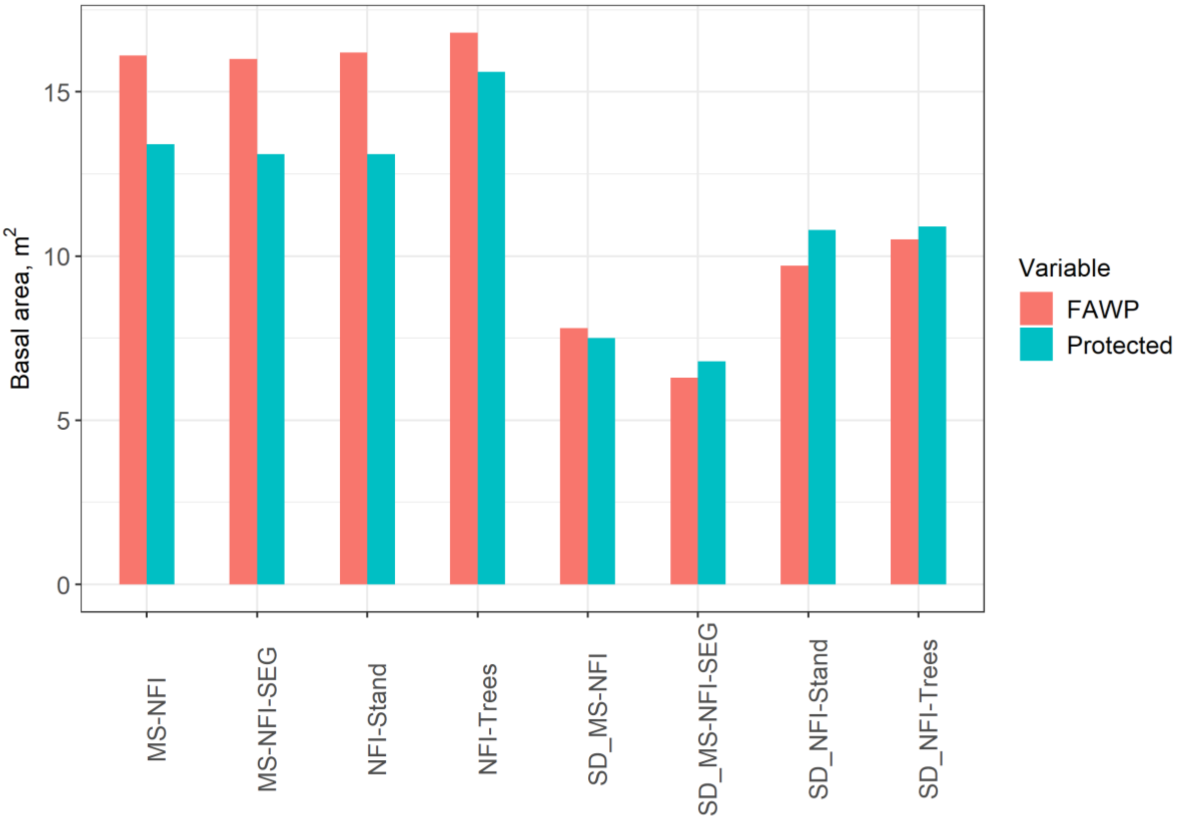


**Figure S3.** Basal areas and corresponding standard deviations on forested land available for wood production (FAWP) and on protected areas. Datasets are NFI trees measured on the plot and stand around the plot, MS-NFI and segmented MS-NFI.
